# Zebrafish models for human ALA-dehydratase-deficient porphyria (ADP) and hereditary coproporphyria (HCP) generated with TALEN and CRISPR-Cas9

**DOI:** 10.1101/109553

**Authors:** Shuqing Zhang, Jiao Meng, Zhijie Niu, Yikai Huang, Jingjing Wang, Xiong Su, Yi Zhou, Han Wang

**Author notes:** To whom correspondence should be addressed at: Han Wang, Center for Circadian Clocks, Soochow University, 199 Renai Road, Suzhou, Jiangsu 215123, China.; or.

## Abstract

Defects in the enzymes involved in heme biosynthesis result in a group of human metabolic genetic disorders known as porphyrias. Using a zebrafish model for human hepatoerythropoietic porphyria (HEP), caused by defective uroporphyrinogen decarboxylase (Urod), the fifth enzyme in the heme biosynthesis pathway, we recently have found a novel aspect of porphyria pathogenesis. However, no hereditable zebrafish models with genetic mutations of *alad* and *cpox*, encoding the second enzyme delta-aminolevulinate dehydratase (Alad) and the sixth enzyme coproporphyrinogen oxidase (Cpox), have been established to date. Here we employed site-specific genome-editing tools transcription activator-like effector nuclease (TALEN) and clustered regularly interspaced short palindromic repeats (CRISPR)/CRISPR-associated protein 9 (Cas9) to generate zebrafish mutants for *alad* and *cpox*. These zebrafish mutants display phenotypes of heme deficiency, hypochromia, abnormal erythrocytic maturation and accumulation of heme precursor intermediates, reminiscent of human ALA-dehydratase-deficient porphyria (ADP) and hereditary coproporphyrian (HCP), respectively. Further, we observed altered expression of genes involved in heme biosynthesis and degradation and particularly down-regulation of exocrine pancreatic zymogens in ADP (*alad-/-*) and HCP (*cpox-/-*) fishes. These two zebrafish porphyria models can survive at least 7 days and thus provide invaluable resources for elucidating novel pathological aspects of porphyrias, evaluating mutated forms of human *ALAD* and *CPOX*, discovering new therapeutic targets and developing effective drugs for these complex genetic diseases. Our studies also highlight generation of zebrafish models for human diseases with two versatile genome-editing tools.

## Introduction

The porphyrias are a group of rare and clinically complex metabolic disorders, each caused by genetic mutations that result in defective enzymes involved in eight highly conserved reactions of heme biosynthesis (Balwani and Desnick, 2012; Puy et al., 2010). Deficiencies of the second enzyme delta-aminolevulinate dehydratase (ALAD; EC 4.2.1.24) in the heme biosynthesis pathway cause autosomal recessive ALA-dehydratase-deficient porphyria (ADP; Online Mendelian Inheritance in Man, OMIM 125270, John Hopkins University, Baltimore, MD) (Jaffe and Stith, 2007; Yano and Kondo, 1998); whereas deficiencies of the sixth enzyme coproporphyrinogen oxidase (CPOX, EC 1.3.3.3) lead to autosomal dominant hereditary coproporphyria (HCP; OMIM 121300) (Whatley et al., 2009). Both porphyrias are acute hepatic and clinically manifest neurologic attacks, abdominal pain, nausea, vomiting, tachycardia, and hypertension (Siegesmund et al., 2010). The diagnosis of the two porphyrias is primarily through detection of elevated levels of delta-aminolevulinic acid (ALA), coproporphyrin III and porphobilinogen (PBG) in urine, feces and blood, combined with enzymatic assays and mutation analysis (Balwani and Desnick, 2012; Cappellini et al., 2010; Maruno et al., 2001; Puy et al., 2010).

The zebrafish (*Danio rerio*) has been a model for investigating molecular genetic mechanisms underlying numerous human disorders including hematopoietic diseases (Dooley and Zon, 2000; Lieschke and Currie, 2007), and also has been particularly effective for high-throughput drug screens in whole organism (MacRae and Peterson, 2015; North et al., 2007; Poureetezadi et al., 2014; Rennekamp and Peterson, 2015; Zon and Peterson, 2005). Using a zebrafish model for human hepatoerythropoietic porphyria (HEP; OMIM 176100) (Wang et al., 1998), we recently have uncovered a novel aspect of porphyric pathogenesis, i.e., heme regulates exocrine pancreatic zymogens through the Bach1b/Nrf2a-MafK pathway (Wang et al., 2007; Zhang et al., 2014), demonstrating that the utilities and power of zebrafish disease models for delineating novel pathogenesis mechanisms (Santoriello and Zon, 2012). However, no hereditable zebrafish *alad* and *cpox* models with genetic mutations have been generated to date. A medaka (*Oryzias latipes*) *whiteout* (*who*) mutant was previously characterized as a model for human ADP based upon its phenotype of hypochromic anemia (Sakamoto et al., 2004), but medaka spawns relative fewer eggs (20~40) each time (Furutani-Seiki and Wittbrodt, 2004) and has not yet been developed for large-scale drug screens. Even though *cpox*-Morpholino-injected zebrafish embryos were reported as an *in vivo* assay system for assessing biological activities of human *CPOX* mutations (Hanaoka et al., 2006), these *cpox* morphants are not genetically stable and particularly difficult to scale up for drug discoveries. A Nakano mouse displays reduced CPOX enzymatic activities resulted from a *Cpox* mutation (Mori et al., 2013), however, the Nakano mouse also harbors another mutation in the *Crybg3* gene tightly linked with *Cpox* on mouse chromosome 16. Thus the Nakano mouse also manifests cataracts that human HCP patients do not develop, whereby compounding its phenotype and limiting its use in investigating novel aspects of porphyria pathogenesis. Hence, zebrafish *alad* and *cpox* genetic models are needed.

Transcription activator-like effector nuclease (TALEN) and clustered regularly interspaced short palindromic repeats (CRISPR)/CRISPR-associated protein 9 (Cas9) are two site-specific genome-editing systems, each composed of a DNA-recognizing component and a DNA-cleavage component (Hwang et al., 2013; Irion et al., 2014; Wei et al., 2013). In the TALEN system, the two TALE domains are the DNA-recognizing component and the two FokI domains are the DNA-cleavage component (Huang et al., 2014), while in the CRISPR-Cas9 system, the single guide-RNA (gRNA) is in charge of DNA-recognizing and the Cas9 endonuclease in charge of DNA-cleavage (Hsu et al., 2014). A pair of TALENs or one gRNA along with the Cas9 protein suffice to generate site-specific DNA double-strand breaks (DSBs) that trigger the endogenous nonhomologous end joining (NHEJ) DNA repair pathway to induce indel mutations in targeted genes in numerous species including zebrafish (Cong and Zhang, 2015; Wei et al., 2013).

Here we used TALEN and CRISPR-Cas9 to generate zebrafish mutants for *alad* and *cpox.* Characterization of these two zebrafish mutants found that they both manifest phenotypes of heme deficiency, hypochromia, abnormal erythrocytic maturation and accumulation of heme precursor intermediates, and in particular that *cpox* mutants display reddish autofluorescence, resembling human ALA-dehydratase-deficient porphyria (ADP) and hereditary coproporphyrian (HCP), respectively. These zebrafish new models of human porphyrias are invaluable for elucidating novel pathological aspects of porphyrias, in vivo assessments of mutations of these human disease genes, discovering new therapeutic targets and developing effective drugs for these complex genetic diseases.

## Results

### Generation of zebrafish *alad* mutants with TALEN

A pair of TALENs was designed to target exon 3 of zebrafish *alad* (ENSDARG00000052815) with a BsmA I restriction endonuclease site in the targeted fragment for subsequent mutant identification (Fig. 1A). Following microinjection of this pair of *alad* TALEN mRNAs into single-cell zebrafish embryos, we extracted DNAs from injected and control embryos. By PCR amplification and restriction endonuclease digestion analysis, we estimated a mutagenesis rate of approximately 44% in F_0_ embryos (Fig. S1A). To identify the induced mutation types, the uncleaved DNA fragments were then cloned and sequenced. Sequencing results showed that there are several types of deletions in the targeted fragment (Fig. 1B) (Xiao et al., 2013). The remaining F_0_ fish were raised to adulthood and then crossed with wild-type fish to produce F_1_. To screen for inheritable mutations, DNAs extracted from approximately 10 F_1_ embryos of each cross were used for PCR amplification and enzymatic digestion analysis. The germline transmission efficiency of *alad* TALEN-induced mutations was estimated to be approximately 20% (Supplementary Table S1). Siblings of the F_1_ fish carrying inheritable mutations were raised to adulthood and re-identified by PCR, restriction endonuclease digestion and sequencing of fin-clipped DNAs. An *alad* mutant line carrying a 14-bp deletion resulting in a frame shift starting the 51th amino acid (ZFIN allele nomenclature designation, *alad^sus003^*) (Fig. 1C) was used for the further experiments.

**Fig. 1.**
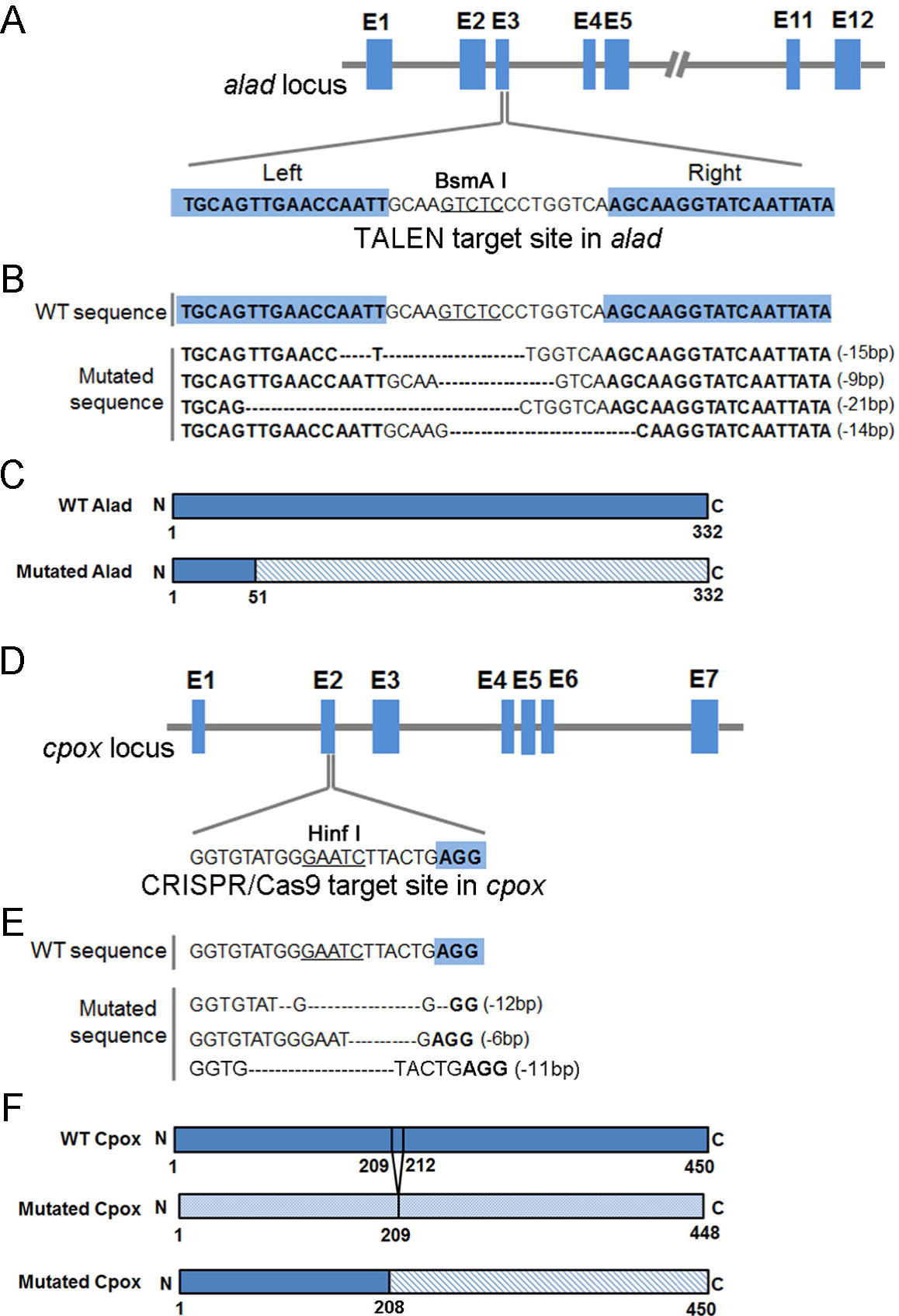
TALEN and CRISPR-Cas9 targeted sites and mutated sequences. (A) TALENs were designed to target the third exon of the zebrafish *alad* gene. The wild-type (WT) sequence with the two TALEN arms is highlighted in blue, and the BsmA I restriction endonuclease site for genotyping is underlined. (B) WT sequence and F_0_ mutated sequence with indels at the *alad* TALEN-targeted fragment. (C) A *alad* mutant line contains a 14-bp deletion resulting in a frame shift starting at the 51^st^ amino acid. (D) A 20-nt gRNA was designed to target the second exon of the zebrafish *cpox* gene. The WT sequence with the PAM is highlighted in blue, and the Hinf I restriction endonuclease site for genotyping is underlined. (E) WT sequence and F_0_ mutated sequences with indels at the *cpox* CRISPR-Cas9-targeted fragment. (F) Two *cpox* mutant lines, one containing a 6-bp deletion resulting in deletion of the 2 amino acids (210 Leu and 211 Thr) and the other having a 11-bp delection resulting a truncated peptide with only 208 amino acids.

### Generation of zebrafish *cpox* mutants with CRISPR-Cas9

To generate zebrafish mutants for *cpox* (ENSDARG00000062025), we selected a 20-nt gRNA immediate to a PAM (protospacer adjacent motif) sequence (NGG) located at exon 2 of *cpox*, and the targeted fragment also contains a Hinf I restriction endonuclease site for subsequent genotyping (Fig. 1D). Single-cell zebrafish embryos were microinjected with the synthetic Cas9 mRNA and *cpox* gRNA. DNAs extracted from injected and control embryos were PCR amplified. Restriction endonuclease digestion of the PCR products detected approximately 44% to 49% of mutagenesis frequency induced by gRNA-Cas9 (Fig. S1B). Cloning and sequencing of the uncleaved DNA fragments showed several types of small deletions adjacent to PAM in F_0_ embryos (Fig. 1E). The remaining F_0_ embryos were raised to adulthood. DNAs extracted from the F_1_ embryos produced by crossing F_0_ fish with wild-type fish were PCR amplified. Restriction endonuclease digestion of the PCR products identified the corresponding F_0_ fish carrying inheritable mutations. The germline transmission efficiency of the *cpox* CRISPR/Cas9-induced mutations was approximately 17% (Supplementary Table S1). Siblings of the F_1_ fish identified as carrying inheritable mutations were raised to adulthood and re-identified by PCR, restriction endonuclease digestion and sequencing of fin-clipped DNAs. Two *cpox* mutant lines, one harboring a 6-bp deletion resulting in deletion of 2 amino acids (ZFIN allele nomenclature designation, *cpox^sus004^*) in a highly conserved region (Hanaoka et al., 2006) and the other possessing a 11-bp deletion resulting in a truncated peptide with only 208 amino acids (ZFIN allele nomenclature designation, *cpox^sus005^*), were identified (Fig. 1F). These two *cpox* mutant fishes display similar phenotypes and were used in the following experiments.

### Zebrafish *alad-/-* and *cpox-/-* mutants display severe hypochromic anemia

The *alad-/-* and *cpox-/-* mutant embryos exhibit hypochromic phenotype as early as 48 hours post fertilization (hpf) (Fig. 2A). The circulating blood cells of the wild-type pericardia are reddish, while those of the *alad-/-* and *cpox-/-* mutant pericardia are pale (Fig. 2A). In addition, the *cpox-/-* mutant displays reddish autofluorescence (see the result below), presumably due to the excessive accumulation of photosensitive porphyrin. Although heterozygous *alad+/-* and *cpox+/-* fish appear unaffected outwardly and can grow to adulthood, homozygous embryos die due to severe embryonic anemia. The mass death of *alad-/-* homozygotes starts at 7 dpf, and their survival rate drops to 0% at 14 dpf (Fig. 2D), suggesting that *alad-/-* homozygous larvae cannot survive longer than 13 days. The mass death of *cpox-/-* starts at 6 dpf, and *cpox-/-* homozygous larvae cannot survive longer than 11 days (Fig. 2D). Homozygous *alad-/-* and *cpox-/-* larvae that survive past day 10 (Fig. 2B) are pale, and have much shorter body length (Fig. 2C) but do not have observable ingested food in the intestine compared to wild-type control larvae (Fig. 2B).

**Fig. 2.**
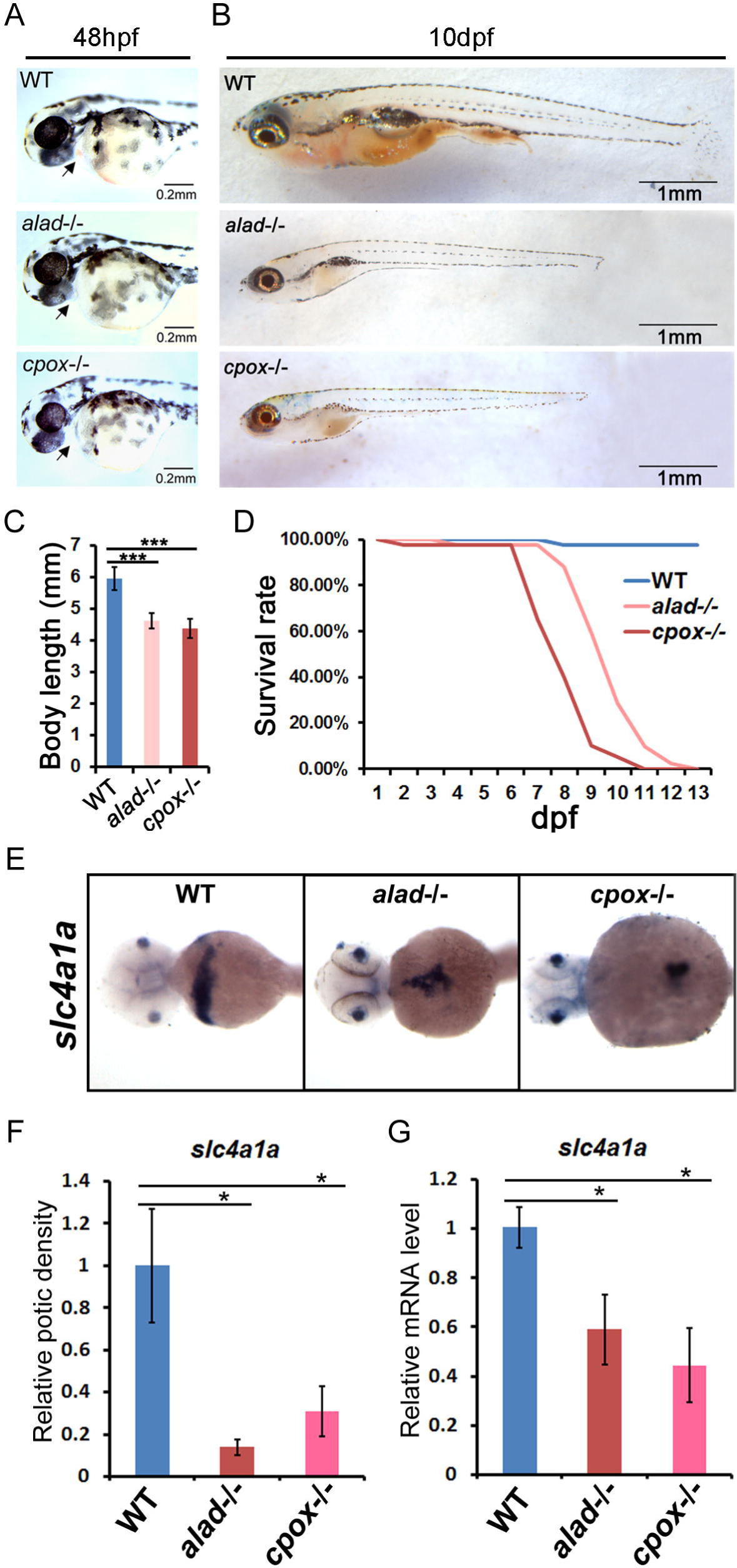
Phenotypes of homozygous mutants with mutated alad and cpox genes. (A) Lateral views of WT, *alad-/-* and *cpox-/-* (6-bp deletion) zebrafish embryos at 48 hpf. Arrows indicate the pericardium. (B) Lateral views of WT, *alad-/-* and *cpox-/-* (6-bp deletion) zebrafish larvae at 10 dpf. (C) Body lengths of *alad-/-* and *cpox-/-* (6-bp deletion) larvae at 10 dpf compared with wild types. 25 larvae per group were measured for statistical analysis. Student’s *t*-tests were conducted. ***P<0.001. (D) Survival rate of *alad-/-* and *cpox-/-* (6-bp deletion) from 1 dpf to 13d pf compared with wild types. 40 larvae per group were examined for statistical analysis. (E) *In situ* hybridization staining shows the down-regulation of *slc4a1a* in *alad-/-* and *cpox-/-* (11-bp deletion) embryos at 48 hpf. (F) Mean optic densities of *in situ* hybridization staining of a group of larvae (10-12 each) corresponding to Fig. 2E were quantified by using ImageJ. Student’s *t*-tests were conducted. *P<0.05. (G) qRT-PCR analysis shows down-regulation of *slc4a1a* in *alad-/-* and *cpox-/-* (11-bp deletion). Student’s *t*-tests were conducted. *P<0.05.

### Abnormal erythrocytic maturation in *alad-/-* and *cpox-/-* mutant larvae

Whole-mount *in situ* hybridization and qRT-PCR show the expression of *slc4a1a*, a terminal erythroid marker gene, significantly down regulated in *alad-/-* and *cpox-/-* mutants (Fig. 2E, F and G), suggesting insufficient erythroid cells in these two porphyric mutant zebrafish. Whole-embryo *o*-dianisidine staining also revealed that the hemoglobin contents of blood cells are severely reduced in *alad-/-* and *cpox-/-* mutants compared with wild types. Hemoglobin of wild-type embryos is distributed widely in the ventral part of the yolk from 32hpf to 72hpf (Fig. 3A, D and G), while that of *alad-/-* (Fig. 3B, E and H) and *cpox-/-* (Fig. 3C, F and I) mutant embryos can be observed only in a small and narrow area and is markedly reduced compared with wild types (Fig. 3J). We also performed the hematoxylin-eosin (HE) staining after *o*-dianisidine staining. Results show that blood cells in *alad-/-* and *cpox-/-* lack the red color of *o*-dianisidine staining signal compared with wild-type blood cells, suggesting loss of hemoglobin in these two porphyric mutant zebrafish (Fig. S2A). The blood cell density in section slides is also significantly reduced in *alad-/-* and *cpox-/-* (Fig. S2B), suggesting the number of blood cells is significantly decreased in *alad-/-* and *cpox-/-*.

Wright staining of circulating embryonic red blood cells showed absence of acidophilic materials (Fig. 3L-T), especially the cytoplasm in *alad-/-* does not show any red color at 108 hpf (Fig. 3S). Blood smears assay showed that blood cells of *alad-/-* and *cpox-/-* mutants have larger nuclei compared with wild-type cells at 72 hpf and 108 hpf (Fig. 3U), suggesting that the erythrocytes of the two mutants differentiate abnormally.

Furthermore, heme contents of the circulating embryonic red blood cells are significantly reduced in *alad-/-* and *cpox-/-* mutants than wild types (Fig. 3V), and the mRNA levels of *hbae1* and *hbbe2*, encoding α_e1_-globin and β_e2_-globin, are significantly down-regulated in *alad-/-* and *cpox-/-* mutant fishes (Fig. 3K), indicating that heme biosynthesis is disrupted in *alad-/-* and *cpox-/-* mutants, and subsequently leading to the reduced hemoglobin (Tahara et al., 2004a; Tahara et al., 2004b) and abnormal erythrocyte differentiation.

**Fig. 3.**
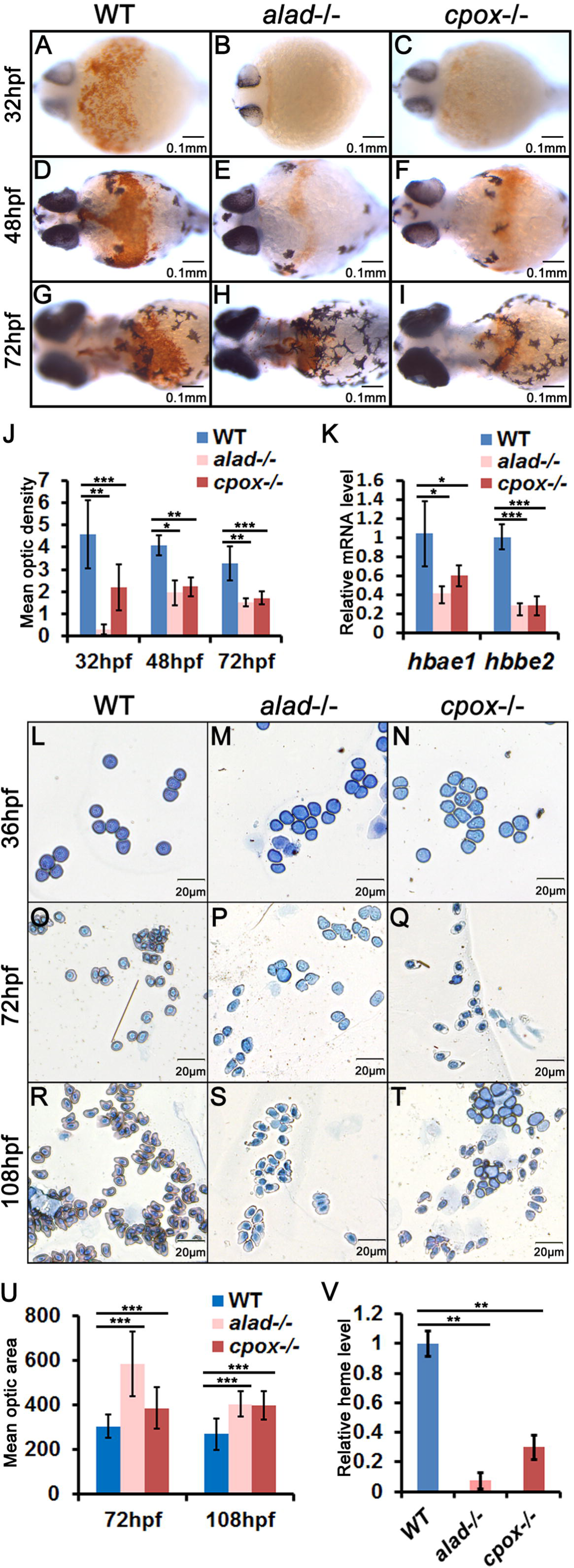
Abnormality of blood cells in *alad-/-* and *cpxo-/-* mutants. Whole-mount *o*-dianisidine staining of WT (A, D and G), *alad-/-* (B, E and H) and *cpxo*-/- (6-bp deletion) (C, F and I) embryos at 32 hpf (A, B and C), 48 hpf (D, E and F), and 72 hpf (G, H and I). Shown are ventral views. (J) Statistical analysis of optic densities of whole-mount *o*-dianisidine staining, shown in A, B, C, D, E, F, G, H and I, measured by ImageJ. 20 embryos per group were measured for statistical analysis. Student’s *t*-tests were conducted. *P<0.05, **P<0.01, ***P<0.001. (K) Relative mRNA levels of *hbae1* and *hbbe2* in WT, *alad-/-* and *cpxo-/-* (6-bp deletion) embryos at 48hpf. Circulating red blood cells of WT (L, O and R), *alad-/-* (M, P and S) and *cpxo*-/- (6-bp deletion) (N, Q and T) embryos/larvae at 36 hpf (L, M and N), 72 hpf (O, P and Q), and 108 hpf (R, S and T) with Wright staining. (U) Statistical analysis of optic nuclear areas shown in L, M, N, O, P, Q, R, S, and T. The nuclei of *alad-/-* and *cpox-/-* (6-bp deletion) are larger than those of WT at 72 hpf and 108 hpf. Optic nuclear areas were measured by ImageJ. 30 blood cells per group were measured for statistical analysis. Student’s *t*-tests were conducted. ***P<0.001. (V) Relative heme levels of circulating red blood cells in WT, *alad-/-* and *cpxo-/-* (6-bp deletion) embryos at 7 dpf, determined by the QuantiChrom^TM^ Heme Assay Kit.

### Homozygous *alad-/-* and *cpox-/-* fishes resemble ADP and HCP

The substrate of aminolevulinic acid dehydrase (Alad) is porphyrin precursor δ-aminolevulinic acid (ALA), which normally does not accumulate in significant amounts. The hepatic porphyrias are characterized by overproduction and accumulation of ALA (Cappellini et al., 2010; Costa et al., 1997; Puy et al., 2010; Yano and Kondo, 1998). HPLC (high-performance liquid chromatography) assays showed excessive amounts of ALA in *alad-/-* embryos compared with wild-type embryos (Fig. 4A), indicative of defective Alad in the *alad-/-* mutant. In *cpox-/-*, the reddish autofluorescence of the blood cells can be observed in the yolk as early as 30hpf (Fig. 4C, D), and HPLC assays also showed excessive amounts coproporphyrinogen III (CP), the substrate of coproporphyrinogen oxidase (Cpox), in *copx-/-* embryos compared with wild types, indicating the excessive accumulation of photosensitive coproporphyrinogen III of the heme biosynthesis pathway.

**Fig. 4.**
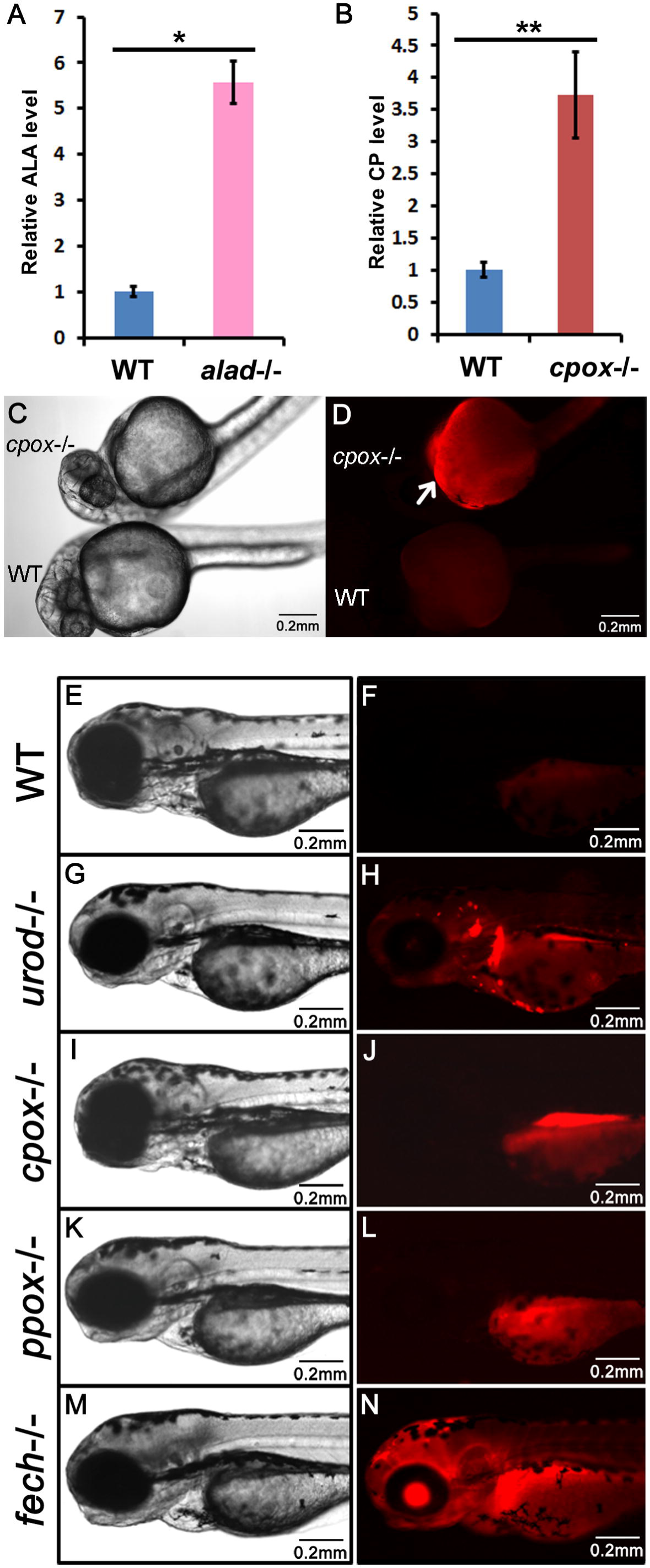
Accumulation of metabolic intermediates in *alad-/-* and *cpxo-/-* mutants. (A) ALA measurements with HPLC (High-performance liquid chromatography) in WT and *alad-/-* larvae at 7 dpf. Relative fluorescence of ALA peak in WT and *alad-/-*. (B) CP measurements with HPLC in WT and *cpox-/-* (11-bp deletion) larvae at 7 dpf. Relative fluorescence of CP peak in WT and *cpox-/-* (11-bp deletion). Reddish autofluorescence of *cpox-/-* embryo at 32 hpf (C, D), the top embryo is *cpox-/-* mutants, the bottom one is the wild-type siblings. (C) Bright light image. (D) Image under a fluorescent microscope with a rhodamine filter. The arrow indicates autofluorescence of the yolk sac of the *cpox-/-* (6-bp deletion) embryo. Autofluorescence analysis of larvae of wild-type (E, F), *urod-/-* (G, H), *cpox-/-* (6-bp deletion) (I, J), *ppox-/-* (L, L) and *fech-/-* (M, N) at 72hpf. (E, G, I, K and M), Bright light image. (F, H, J, L and N) Image under a fluorescent microscope with a rhodamine filter.

To determine whether *alad* mRNA or *cpox* mRNA can rescue the phenotypes of *alad-/-* or *cpox-/-* mutant fishes, we microinjected theses mRNAs into the mutant and wild-type control embryos (Fig. 5). *o*-Dianisidine staining showed that the hemoglobin content was increased to 60% of the normal level in *alald-/-* and 80% in *cpox-/-* following mRNA microinjection, indicating that the hypochromic anemia phenotype of *alald-/-* and *cpox-/-* is resulted from the mutated *alad* or *cpox* genes. Taken together, homozygous *alad-/-* and *cpox-/-* fishes display phenotypes of hypochromia, heme deficiency, abnormal erythrocytic maturation and accumulation of heme precursor intermediates, mimicking human ALA-dehydratase-deficient porphyria (ADP) and hereditary coproporphyrian (HCP).

**Fig. 5.**
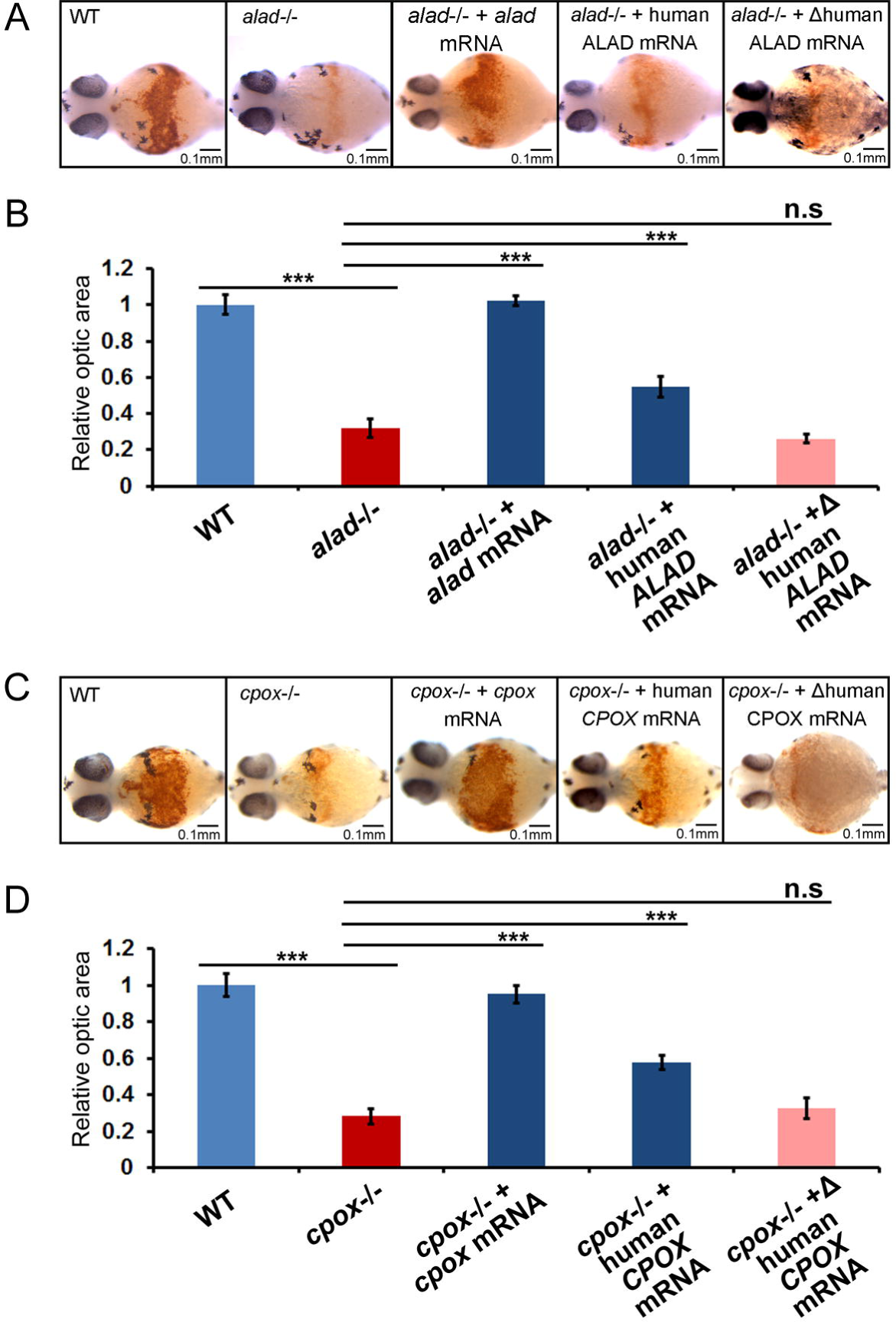
Rescuing the phenotypes of zebrafish *alad-/-* and *cpxo-/-* mutants by zebrafish and human genes. (A) Whole-mount *o*-dianisidine staining of wild-type embryos, *alad-/-* embryos, *alad-/-* embryos injected with zebrafish *alad* mRNA and *alad-/-* embryos injected with human *ALAD* mRNA at 48 hpf. (B) Statistical analysis of optic densities measured by ImageJ. 25 embryos per group were measured for statistical analysis. Student’s *t*-tests were conducted. *P<0.05, ***P<0.001. (C) Whole-mount *o*-dianisidine staining of wild-type embryos, *cpox-/-* (6-bp deletion)embryos, *cpox-/-* (6-bp deletion) embryos injected with zebrafish *cpox* mRNA and *cpox-/-* embryos injected with human *CPOX* mRNA at 48 hpf. (D) Statistical analysis of optic blood cells about C. Optic densities were measured by ImageJ, 20 embryos per group were measured for statistical analysis. Student’s *t*-tests were conducted. *P<0.05, ***P<0.001.

### Evaluating the mutated forms of human ALAD and CPOX in homozygous *alad-/-* and *cpox-/-* fishes

The zebrafish Alad and Cpox proteins are highly similar in sequence to human and mouse counterparts (Fig. S3) (Hanaoka et al., 2006). In order to examine whether the biological activities of these two enzymes are conserved between zebrafish and human, *alad-/-* and *cpox-/-* embryos were microinjected with human *ALAD* or *CPOX* mRNAs at one-cell stage, respectively. *o*-Dianisidine staining showed that hemoglobin was significantly increased after injection compared with the mutant embryos without injection at 48hpf (Fig. 5), indicating that the hypochromic anemia phenotype of *alad-/-* and *cpox-/-* also can be rescued by human *ALAD* and *CPOX*. In contrast, microinjected with mutated forms of human *ALAD* or *CPOX* mRNAs from the ADP and HCP patients (Ishida et al., 1992; Rosipal et al., 1999) did not show any rescue effect (Fig. 5). These results suggest that the functions of these two enzymes are conserved between zebrafish and human, and we are able to effectively use homozygous *alad-/-* and *cpox-/-* fishes to evaluate mutations of these two human disease genes.

### Expression patterns of genes involved in heme biosynthesis and degradation, and exocrine pancreatic zymogens are disrupted in ADP (*alad-/-*) and HCP (*cpox-/-*) fishes

The heme biosynthesis pathway is tightly regulated, and in particular heme negatively regulates the first and rate-limiting enzyme Alas1 in the pathway in non-erythroid cells (Furuyama et al., 2007). qRT-PCR analysis showed that *alas1* is up-regulated in both ADP and HCP fishes (Fig. 6A, B), likely resulted from heme deficiencies caused by *alad* and *cpox* mutations. In contrast, heme can enhance its own synthesis in erythroid cells by up-regulating *alas2* (Chiabrando et al., 2014), and we observed two *hmbs* (encoding the third enzyme Hmbs in the heme biosynthesis pathway) genes, *hmbsa* and *hmbsb* in zebrafish (data not shown). While *alas2* and *hmbsa* are up-regulated but *alad* itself and *hmbsb* are down-regulated in ADP larvae (Fig. 6A), all genes including upstream and downstream of *cpox* except *ppox* are up-regulated in HCP larvae (Fig. 6B).

**Fig. 6.**
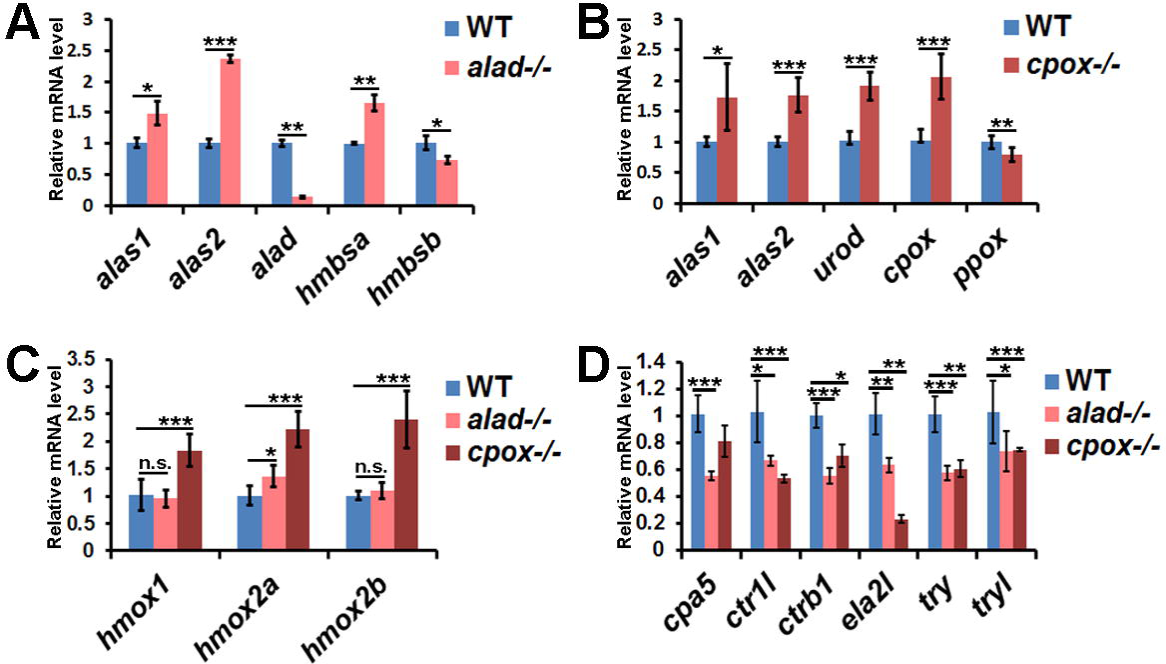
Altered expression of genes involved in heme biosynthesis and degradation, and exocrine pancreatic zymogens in *alad-/-* and *cpox-/-* mutants detected by qRT-PCR analysis. (A) Altered expression of genes involved in heme biosynthesis including *alas1*, *alas2*, *alad*, *hmbsa* and *hmsb* in *alald-/-* at 48 hpf. (B) Altered expression of genes involved in heme biosynthesis including *alas1*, *alas2*, *urod*, *cpox* and *ppox* in *cpox-/-* (6-bp deletion) at 48 hpf. (C) Altered expression of genes involved in heme degradation including *hmox1*, *hmox2a* and *hmox2b* in *alald-/-* and *cpox-/-* (6-bp deletion) at 48 hpf. (D) Down-regulation of exocrine pancreatic zymogens in *alald-/-* and *cpox-/-* (6-bp deletion) at 72 hpf. Student’s *t*-tests were conducted. *P<0.05, **P<0.01, ***P<0.001, n.s., not significant.

Heme oxygenase (Hmox) is the rate-limiting enzyme of the heme degradation pathway, and can be induced by heme and the oxidative stress of organisms (Applegate et al., 1991; Igarashi and Sun, 2006). Zebrafish contain three *hmox* genes, *hmox1*, *hmox2a* and *hmox2b*. While *hmox1*, *hmox2a* and *hmox2b* are up-regulated in HCP larvae, only *hmox2a* is up-regulated in ADP larvae (Fig. 6C), indicating high levels of the oxidative stress in these two mutants especially in HCP larvae, likely due to accumulation of intermediate products of the heme biosynthesis pathway.

Our previous studies found that heme deficiencies result in down-regulation of six peptidase precursor genes in the zebrafish exocrine pancreas (Wang et al., 2007; Zhang et al., 2014). Here qRT-PCR analysis showed that all the six zymogens are also down-regulated in both ADP and HCP larvae (Fig. 6D), providing strong support for our previous finding (Wang et al., 2007; Zhang et al., 2014).

### Complex syndromes in different zebrafish porphyria models

Porphyrias show a group of complex syndromes, characteristic of diverse clinical manifestations (Balwani and Desnick, 2012; Cappellini et al., 2010; Puy et al., 2010). We also observed diverse phenotypes in the four zebrafish porphyria models with defective Urod (Wang et al., 1998), Cpox (this study), Ppox (Dooley et al., 2008) and Fech (Childs et al., 2000). First, different autofluorescence patterns can be observed in these four zebrafish heme-deficient models (Fig. 4E-4N). Reddish fluorescence displays primarily in blood cells and internal organs of HEP (*urod-/-*) larvae (Fig. 4H), primarily in the internal organs and yolk of HCP (*cpox-/-*) (Fig. 4J) and VP (*ppox-/-*) larvae (Fig. 4L), and in the whole body, especially in the lens, brain and liver of EPP (*fech-/-*) larvae (Fig. 4N). Second, light-induced pericardial edemata have been observed in HEP (*urod-/-*) larvae but not other heme-deficient zebrafish models. As shown in Fig. S4, the appearance of HEP (*urod-/-*) larvae protected under dark condition are normal, while HEP (*urod-/-*) larvae under light exposure display an exacerbating pericardial edema phenotype from 72 hpf to 144 hpf, which cannot be observed in HCP (*cpox-/-*) larvae. The distinct patterns of reddish fluorescence and light-induced pericardial edemata may be resulted from accumulation of different porphyrins or porphyrin precursors, and these mutants provide tools for reveal the complex pathological mechanisms of human porphyrias.

## Discussion

Zebrafish have become a bona fide vertebrate model for studying numerous human diseases (Dooley and Zon, 2000; Lieschke and Currie, 2007). Heme biosynthesis is conserved throughout vertebrate evolution (Detrich et al., 1995), all 8 of human genes involved in heme biosynthesis pathway have zebrafish orthologues (Tzou et al., 2014). Four zebrafish mutants *sauternes/alas2* (Brownlie et al., 1998), *yquem/urod* (Wang et al., 1998), *montalcino/ppox* (Dooley et al., 2008) and *dracula/fech* (Childs et al., 2000) were identified in large-scale forward genetic mutagenesis screens (Ransom et al., 1996; Weinstein et al., 1996). A knock-in mouse carrying human *UROS* P248Q missense mutation (*Uros*^mut248^) represents a model of the congenital erythropoietic porphyria (CEP) (Ged et al., 2006), *Pbgd*-deficient mice generated by gene targeting was reported to exhibit the typical biochemical characteristics of human acute intermittent porphyria (AIP) (Ged et al., 2006; Lindberg et al., 1996), a *URO-D*^+/-^ mouse was generated by homologous recombination as a model of familial porphyria cutanea tarda (PCT) (Phillips et al., 2001), a South African variegate porphyria (VP) mouse model was generated by knocking in the human R57W mutation (Medlock et al., 2002; Meissner et al., 1996), and a BALB/c *Fech*^m1Pas^ mouse was generated by a mutagenesis screen using ethylnitrosourea (ENU) as a model of erythropoietic protoporphyria (EPP) (Tutois et al., 1991)(Table 1). Nevertheless, zebrafish models for ALA-dehydratase deficient porphyria (ADP), acute intermittent porphyria (AIP), congenital erythropoietic porphyria (CEP) and hereditary coproporphyria (HCP) have not been reported to date. Here, we generated two zebrafish models for ADP and HCP with TALEN and CRISPR-Cas9, respectively. Our work emphasizes that the engineered endonucleases (EENs) TALEN and CRISPR-Cas9 are efficient tools to generate mutations in zebrafish as models for heritable human disease.

**Table 1.**
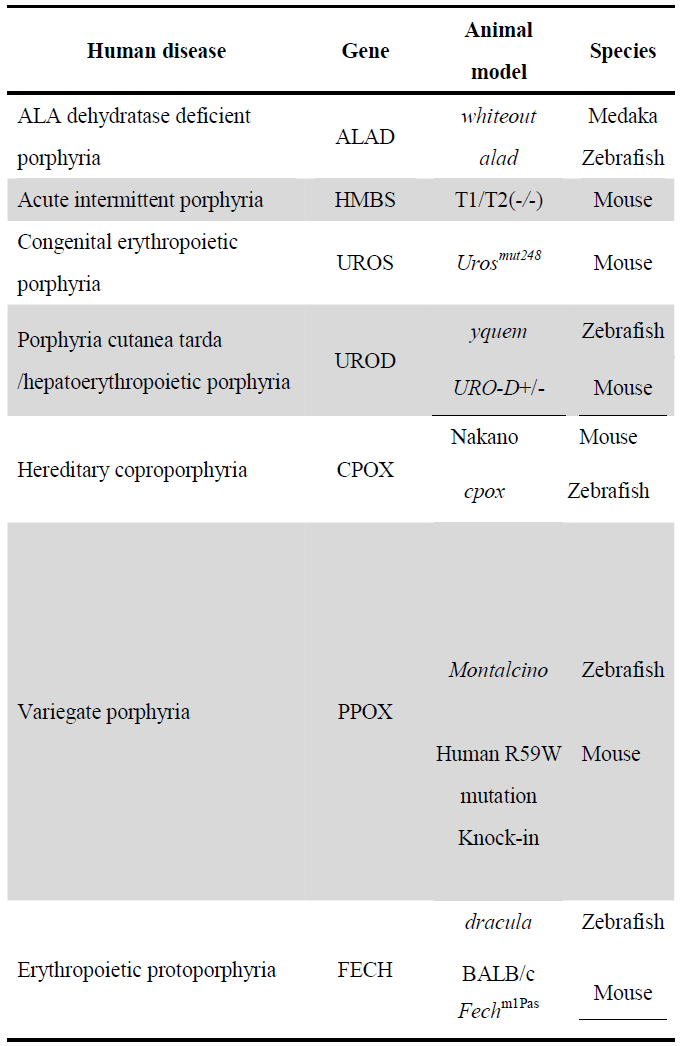
Animal models for human porphyrias

Our studies of *alad-/-* and *cpox-/-* mutants demonstrate that defects in Alad or Cpox cause hypochromic anemia and abnormal erythrocytic maturation in zebrafish. The accumulation of ALA and CP, the metabolic intermediates, in the heme synthetic pathway in *alad-/-* and *cpox-/-* mutants (Fig. 4A, B) indicates that Alad and Cpox enzymatic activities are impaired by these mutations. Expression of the genes involved in the heme biosynthetic pathway is altered in *alad-/-* and *cpox-/-*, which may be caused by the accumulation of metabolic intermediates and/or heme deficiency. The heme oxygenases participate in the cellular defense from the oxidative damage of reactive oxygen species (ROS) generated by heme precursors (Ryter and Tyrrell, 2000). Although heme levels were reduced in *alad-/-* and *cpox-/-*, the heme-induced heme oxygenase genes were up-regulated likely due to the increased level of oxidative stress. Interestingly, consistent with our previous finding of heme deficiency-induced down-regulation of exocrine pancreatic zymogens (Wang et al., 2007; Zhang et al., 2014), all these six exocrine pancreatic zymogens are significantly down-regulated in ADP (*alad-/-*) and HCP (*cpox-/-*) fishes (Fig. 6D).

Most of the reported human ADP patients are compound heterozygous or heterozygous, with point mutations of *ALAD* gene resulting in amino acid changes and reduced enzyme activity (Ishida et al., 1992; Maruno et al., 2001). For the HCP patients, most are heterozygotes with reduced enzymatic activities and homozygotes are rare (Kuhnel et al., 2000; Lamoril et al., 1995; Nordmann et al., 1983). The enzymatic activities of human ALAD or CPOX were investigated through *in vitro* expression of mutated genes from patients (Martasek et al., 1997; Maruno et al., 2001). Our rescue experiments showed that the biological activities of these two enzymes are conserved between zebrafish and human (Fig. 5), these homozygous zebrafish provided *in vivo* tools for evaluating these human mutated enzymes.

Reddish fluorescence was observed in *cpox-/-* embryos under an epifluorescent stereomicroscope using a rhodamine filter (Fig. 4D). We found that four heme-deficient zebrafish mutants exhibit distinct patterns of autofluorescence in different organs (Fig. 4E-N), indicating the complex syndromes and distinct underlying pathological mechanisms in different types of porphyrias. All eight types of porphyrias have animal models, providing invaluable resources to study the pathogenesis of human porphyrias, and six of them are zebrafish models thus far (Table 1). Circulating blood cells of ADP (*alad-/-*) and HCP (*cpox-/-*) fish lack red color, characteristic of hypochromic anemia in zebrafish, which can be observed as early as 48 hpf. Moreover, both of these two homozygous mutants can live at least 7 days, providing a time window for using these zebrafish models to investigate novel aspects of porphyria pathogenesis.

Zebrafish have been employed as an ideal vertebrate model organism for high-throughout drug screens and development in whole organism (MacRae and Peterson, 2015; North et al., 2007; Rennekamp and Peterson, 2015; Zon and Peterson, 2005). The adult heterozygous zebrafish with deficiencies of *alad* or *cpox* can spawn a large number of embryos each week, homozygous mutant embryos can be easily identified according to conspicuously pale blood cells (Fig. 2, Fig. S4), and the *cpox-/-* homozygous embryos even can be identified as early as 32 hpf according to autofluorescent traits under an epifluorescent stereomicroscope (Fig. 4), which all can facilitate high-throughput chemical screening for discovering of potential drugs for preventing or ameliorating the similar symptoms of ADP and HCP in humans. Potential drugs can be dissolved in the embryo medium and the extent of recovery phenotypes can be easily detected also through *o*-dianisidine staining (Fig. 5), which provide an effective assay for drug screening for human porphyrias using these two zebrafish models.

## Materials and Methods

### Fish husbandry and embryo production

The Soochow University Animal Use and Care Committee approved all animal protocols. Zebrafish wild-type AB strain, and mutant line *alad-/-*, *cpox-/-*, *urod-/-* (Wang et al., 1998), *ppox-/-* (Dooley et al., 2008) and *fech-/-* (Childs et al., 2000) are raised at the Soochow University Zebrafish Facility. Wild-type (WT) and mutant embryos were produced by pair mating, collected for RNA isolation at specified stages. Homozygous mutants were obtained by mating heterozygous fish and then identified under an epifluorescent stereomicroscope (Leica M165 FC). The zebrafish mutant lines *alad* and *cpox* generated in this study (See below) have been deposited in both the China Zebrafish Resource Center (CZRC, http://en.zfish.cn/) and the Zebrafish International Resource Center (ZIRC, http://zebrafish.org/home/guide.php), and the *alad* mutant ZFIN allele nomenclature ID is *alad*^*sus003*^ (14-bp deletion), and the *cpox* ZFIN allele nomenclature mutant IDs are *cpox*^*sus004*^ (6-bp deletion) and *cpox*^*sus005*^ (11-bp deletion), respectively.

### *alad-*TALEN construction and microinjection

The TALEN sites targeting the third exon of zebrafish *alad* were designed using a web-tool TALEN-NT (https://boglab.plp.iastate.edu/) (Doyle et al., 2012). TALEN expression vectors were constructed using the ‘Unit Assembly’ method with Sharkey-AS and Sharkey-R forms of FokI cleavage domains as described previously (Huang et al., 2014), were linearized by NotI and used as templates for TALEN mRNA synthesis with SP6 mMESSAGE mMACHINE Kit (Ambion). TALEN mRNAs encoding each monomer were injected in pair into one-cell zebrafish embryos at concentrations of 300 pg. Primers for amplifying the TALEN-targeted fragment are listed in Supplementary Table S2.

### *cpox*-gRNA construction and microinjection

A 20-nt gRNA was selected to target exon 2 of zebrafish *cpox*. The PCR primers for the generation of the DNA template for *cpox*-gRNA were designed, and a T7 promoter sequence was added to 5’upstream of the gRNA sequence. Cas9 mRNA and gRNA were synthesized as described previously (Hwang et al., 2013). Briefly, the Cas9 mRNA was synthesized by *in vitro* transcription using T7 mMESSAGE mMACHINE Kit (Ambion). The gRNA was *in vitro* transcribed and purified using T7 Riboprobe Systems (Promega). Approximately 300 pg of Cas9 mRNA and 75 pg of gRNA were co-injected into one-cell zebrafish embryos. Primers for amplifying the CRISPR-Cas9 targeted fragment are listed in Supplementary Table S2.

### RNA isolation and quantitative RT (Real Time)-PCR

Total RNAs were extracted from approximately 20-30 larvae with the TRIzol^®^ Reagent according to the manufacturer’s instructions (Invitrogen). The cDNAs synthesized by reverse transcription with the M-MLV reverse transcription kit (Invitrogen) were used as template for quantitative RT-PCR (qRT-PCR) analysis. qRT-PCRs were carried out with the ABI StepOnePlus^TM^ systems, using SYBR^®^ Premix Ex Taq™(TaKaRa) and the following thermal profile: 95^°^C for 3 min, and 40 cycles of 95^°^C, 10 sec; and 58^°^C, 30 sec. Primers are listed in Supplementary Table S2. *actb1* was used as an internal control. Relative mRNA expression levels were quantified using the comparative Ct (ΔCt) method and expressed as 2^-(ΔΔCt)^. Each PCR assay was done with three biological samples.

### Construction of the expression vectors

The *alad* and *cpox* ORF (open reading frame) cDNAs were PCR amplified using zebrafish larvae (120hpf) cDNAs as template, then cloned to pcDNA3.1+, and named *alad*-pcDNA3.1 and *cpox*-pcDNA3.1, respectively. Primers for *alad* and *cpox* ORF cloning are listed in Supplementary Table S2. The human *ALAD* and *CPOX* ORF cDNAs were PCR amplified using the NIH 293T cells’ cDNAs as template, and then cloned to pcDNA3.1+, and named *ALAD*-pcDNA3.1 and *CPOX*-pcDNA3.1, respectively. The mutated ORF cDNAs of human *ALAD* (C^718^-T,)(Ishida et al., 1992) and *CPOX* (G^589^-T)(Rosipal et al., 1999) were generated with Q5^®^ Site-Directed Mutagenesis Kit (NEB) using *ALAD*-pcDNA3.1 and *CPOX*-pcDNA3.1, respectively.

### Rescue experiments and *o*-dianisidine staining

The plasmids of expression vectors *alad*-pcDNA3.1, *cpox*-pcDNA3.1, *ALAD*-pcDNA3.1 and *CPOX*-pcDNA3.1 were linearized, and used as templates for generating capped mRNAs with mMESSAGE mMACHINE^®^ Kit according to the manufacturer’s instructions (Ambion). A mixture of capped mRNA in Tris-HCl (0.01M, pH7.0) were microinjected into one-cell zebrafish embryos. Microinjection controls with the vehicle solution free of mRNAs also were performed. Embryos were collected at 48hpf for the *o*-dianisidine staining. Detection of hemoglobin by *o*-dianisidine was performed as described previously (Ransom et al., 1996). The images were acquired with a stereomicroscope microscope (Leica M165 FC) and a digital camera.

### Blood smears assay, Wright staining and heme level determination

We obtained embryonic blood cells and performed Wright staining as described previously (Brownlie et al., 1998; Hanaoka et al., 2006). Embryos were bled by tail amputation and immediately spread on glass slides, allowed to air dry and stained with wright’s dye for 10 minutes, then washed with 1 X PBS. The images were acquired with a stereomicroscope microscope (Olympus U-HGLGPS) and a digital camera. Circulating blood cells maintained in 1 X PBS were collected from 20 embryos and were counted. The heme level assay was performed with the QuantiChrom^TM^ Heme Assay Kit (BioAssay Systems).

### Whole-mount *in situ* hybridization and HE (hematoxylin-eosin) staining

Whole-mount in situ hybridization experiments were conducted as described previously (Wang et al., 2007). Following the whole-embryo *o*-dianisidine staining, selected embryos were rehydrated in 30% sucrose at 4°C overnight for cryostat sectioning. Frozen embryos were sectioned to 8 μm thickness, and hematoxylin-eosin staining was performed as standard protocol. Sections were photographed using a stereomicroscope (Leica M165 FC) with a digital camera, and the images were analyzed by using ImageJ (National Institutes of Health).

### HPLC (High-performance liquid chromatography) assay

For the ALA assay, approximately 300 larvae each for *alad-/-* and WT control were collected at 7 days postfertilization (dpf) for HPLC assay. These larvae were then promptly rinsed and homogenized in 10 mM HEPES buffer for ALA assay as previously described (Costa et al., 1997). HPLC assay was conducted using 5-Aminolevulinic acid hydrochloride (Sigma) as ALA standard.

For the CP assay, approximately 300 zebrafish larvae at 7 dpf were pulverized into fine powder and homogenized in 1 mL 0.2 M HCl aqueous solution by a tissue grinder. The porphyrins were extracted twice into ether as described (Dailey et al., 2015). The extracts were combined and dried under a stream of nitrogen. Standards and samples were resuspended in 100 μl of ethyl acetate/acetic acid (4/1, v/v). The compounds are separated by liquid chromatography on the Accela LC Systems (Thermo Fisher Scientific, San Jose, CA) using a Hypersil GOLD column (3 μm, 150 × 2.1 mm, Part Number: 25003-152130, Thermo Fisher Scientific). Chromatographic separation was conducted in binary gradient using 0.1% formic acid in water as mobile phase A and 0.1% formic acid in methanol as mobile phase B. The gradient elution program started at 40% B for 5 min, increased to 100% B over 7 min, where it was then held for 6 min before returning to 40% B. The flow rate was 0.2 mL/min. MS analysis of Coproporphyrin III was performed on a TSQ Vantage mass spectrometer (Thermo Fisher Scientific) in positive mode using SRM as described(Fyrestam et al., 2015). Coproprophyrinogen III ( Frontier Scientific) was used as CP standard (Tomokuni et al., 1987).

### Statistical analysis

The data were analyzed with unpaired, two-tailed Student’s *t*-test with Microsoft Excel and the results were shown as mean ± SD. The level of significance was accepted at P<0.05. *, ** and *** represent P<0.05, P<0.01 and P<0.001, respectively.

## ACKNOWLEDGEMENTS

We thank Bo Zhang and Wenbiao Chen for providing vectors for TALEN and CRISPR-Cas9; Yi-Lin Yan and members of our laboratory for helpful comments and suggestions on this study.

## Author contributions

H.W. and S.Z. conceived and designed the experiments. S.Z., J.M., Z.N., and J.W. performed the experiments. Y.H. and X.S. helped measure CP contents. Y.Z. provided reagents. S.Z. and H.W. analyzed the data and wrote the paper.

## Resource Impact

### Clinical issue

Heme is the prosthetic group for a number of important proteins and enzymes such as hemoglobin, catalases, and cytochromes. Heme biosynthesis is catalyzed by a cascade of highly conserved eight enzymatic reactions occurred in the mitochondria and the cytoplasm. Defects in the heme biosynthetic enzymes result in a group of human metabolic genetic disorders known as porphyrias. Using a zebrafish model for human hepatoerythropoietic porphyria (HEP), caused by defective uroporphyrinogen decarboxylase (UROD), the fifth enzyme in the heme biosynthesis pathway, we recently have found a novel aspect of porphyria pathogenesis. However, no hereditable zebrafish models with genetic mutations of *alad* and *cpox*, encoding the second enzyme delta-aminolevulinate dehydratase (ALAD) and the sixth enzyme coproporphyrinogen oxidase (CPOX), have been established to date.

### Results

We employed site-specific genome-editing tools transcription activator-like effector nuclease (TALEN) and clustered regularly interspaced short palindromic repeats (CRISPR)/CRISPR-associated protein 9 (Cas9) to generate zebrafish mutants for *alad* and *cpox*. Characterization of these two zebrafish mutants revealed that they display phenotypes of heme deficiency, hypochromia, abnormal erythrocytic maturation and accumulation of heme precursor intermediates, reminiscent of human ALA-dehydratase-deficient porphyria (ADP) and hereditary coproporphyrian (HCP).

### Implications and future directions

These two zebrafish porphyria models provide invaluable resources for elucidating novel pathological aspects of porphyrias, *in vivo* tools for evaluating human mutated forms of these two enzymes, discovering new therapeutic targets and developing effective drugs for these complex genetic diseases. Our studies also highlight generation of zebrafish models for human diseases with two new versatile genome-editing tools.

